# Dynamin-1 is a potential mediator in Cancer-Related Cognitive Impairment

**DOI:** 10.1101/2024.06.04.597349

**Authors:** Ding Quan Ng, Casey Hudson, Tracy Nguyen, Sukesh Kumar Gupta, Yong Qin Koh, Munjal M. Acharya, Alexandre Chan

**Affiliations:** Department of Clinical Pharmacy Practice, School of Pharmacy & Pharmaceutical Sciences, University of California Irvine, Irvine, CA, USA; Department of Anatomy & Neurobiology, School of Medicine, University of California Irvine, Irvine, CA, USA; Department of Pharmacy, Faculty of Science, National University of Singapore, Singapore; Department of Radiation Oncology, School of Medicine, University of California Irvine, Irvine, CA, USA

**Author notes:** Corresponding author: Alexandre Chan, PharmD, MPH, Chair and Professor of Clinical Pharmacy. Address: 802 W Peltason Dr, Irvine, CA 92697-4625.

**Keywords:** Cancer-related cognitive impairment, chemobrain, dynamin-1, extracellular vesicle, exosome

## Abstract

Dynamin-1 (DNM1) consolidates memory through synaptic transmission and modulation and has been explored as a therapeutic target in Alzheimer’s disease. Through a two-prong approach, this study examined its role in cancer-related cognitive impairment (CRCI) pathogenesis using human and animal models. The human study recruited newly diagnosed, chemotherapy-naïve adolescent and young adult cancer and non-cancer controls to complete a cognitive instrument (FACT-Cog) and blood draws for up to three time points. Concurrently, a syngeneic young-adult WT (C57BL/6 female) mouse model of breast cancer was developed to study DNM1 expression in the brain. Samples from eighty-six participants with 30 adolescent and young adult (AYA) cancer and 56 non-cancer participants were analyzed. DNM1 levels were significantly lower among cancer participants compared to non-cancer prior to treatment. While receiving cancer treatment, cognitively impaired patients were found with a significant downregulation of DNM1, but not among those without impairment. In murine breast cancer-bearing mice receiving chemotherapy, we consistently found a significant decline in DNM1 immunoreactivity in the hippocampal CA1 and CA3 subregions. Observed in both human and animal studies, the downregulation of DNM1 is linked with the onset of CRCI. Future research should explore the potential of DNM1 in CRCI pathogenesis and therapeutics development.

## 1. INTRODUCTION

Cancer-related cognitive impairment (CRCI) is a complication of cancer and associated treatments and encompasses cognitive symptoms including poor concentration and executive functioning^1,2^. Affected patients report challenges with resuming pre-cancer responsibilities at work, school, and home, which markedly reduces their quality of life^3–5^. To date, there remains a lack of effective pharmacological interventions that can be routinely provided to these patients.

Dynamin-1 (DNM1), a protein maximally expressed in the central nervous system^6^, plays a pivotal role in memory formation, and lower DNM1 concentrations have been observed in post-mortem brains of dementia patients^7,8^. As long-term memory impairment is also an important sign of CRCI, it is hypothesized that downregulation in DNM1 may be associated with CRCI pathogenesis.

We previously reported a greater downregulation in DNM1 levels in plasma extracellular vesicles (EV) proteomes of breast cancer survivors with perceived cognitive impairment^9^. The study, however, was limited by the use of pooled participant samples and was thus unable to account for characteristics that could confound the DNM1-CRCI relationship when measuring subjective cognition (e.g. co-occurring psychiatric comorbidities^10^). Another important question is the representativeness of DNM1 levels in plasma EVs with brain activity. It is thus necessary to understand how DNM1 expression changes in the brain with tumor induction and chemotherapy administration to develop a more comprehensive picture of the DNM1-CRCI theory.

This study utilized a two-prong approach to enhance our understanding of the DNM1-CRCI relationship that we observed in our exploratory study^9^. First, we investigated the presence of differential DNM1 expression between cancer patients with and without perceived CRCI (adjusted for confounders) in an existing prospective cohort of newly diagnosed adolescent and young adult (AYA) cancer patients^11^. This was followed by an animal study that compared DNM1 expression in the hippocampus between non-cancer and cancer-bearing mice to evaluate brain-associated changes in DNM1 expression due to cancer and chemotherapy. We hypothesize that DNM1 expressions are significantly downregulated among AYA cancer patients suffering from CRCI and within the hippocampus of cancer-bearing, chemotherapy-treated mice.

## 2. MATERIALS AND METHODS

### 2.1 Human Study

#### 2.1.1 Study design

This study was nested within the ACTS study, a prospective, longitudinal cohort study, conducted in Singapore from 2018 to 2022 (NCT03476070). This cohort was originally created to investigate the incidence of cognitive impairment in AYA cancer patients before, during and after anti-cancer treatment. We received ethics approval from SingHealth Institutional Review Board (CIRB Ref 2017/3139) and all participants provided informed consent before participation. Detailed characteristics are previously described in Chan et al^11^.

#### 2.1.2 Patient eligibility criteria

We recruited two participant groups. The first comprised **AYA cancer patients** who were 15-39 years old and newly diagnosed with cancer. The second group were **non-cancer community controls** who were age-matched to cancer participants, did not have prior cancer diagnosis, and were not immediate family members to any of the cancer participants.

#### 2.1.3 Assessment time points and procedures

Cancer patients were evaluated at three time points (pre-treatment, 3-, and 6-months post-baseline, representing T1, T2 and T3 respectively). Non-cancer controls completed two time points (T1 and T3). All participants completed questionnaires for evaluating perceived cognitive function (FACT-Cog) and neuropsychiatric symptoms (MFSI-SF and RSCL) and provided blood samples at all time points. Sociodemographic and cancer-related characteristics were collected from baseline surveys and medical records^11,12^.

#### 2.1.4 Perceived cognitive decline

Participants with perceived cognitive decline (PCD) were identified based on clinically important difference of ≥10.6-point decline in FACT-Cog total score from baseline^14^.

#### 2.1.5 Plasma collection

9 mL of blood was drawn, stored in ethylenediaminetetraacetic acid tubes, and then centrifuged at 1,069 *× g* for 10 minutes at 4°C by trained personnel. Plasma was aliquoted into cryotubes and stored in a −80°C freezer until analysis.

#### 2.1.6 Extracellular vesicle isolation

Plasma EVs are isolated using differential ultracentrifugation as previously described^9,15^. In brief, the plasma samples collected were diluted with an equal volume of phosphate-buffered saline (PBS) and centrifuged at 2,000 *× g* for 30 min and 12,000□*×*□*g* for 60 minutes, both at 4°C to remove cells and cellular debris. The isolated plasma EVs were obtained by ultracentrifugation at 100,000□*×*□*g* for 18 hours using a Beckman L100-XP Ultracentrifuge and stored at −80°C for further analysis.

#### 2.1.7 Extracellular vesicle visualization

The enriched plasma EVs were prepared for transmission electron microscopy (TEM) visualization by transferring them onto a copper grid (Electron Microscopy Services, USA), washed in water and fixed with 2.5% glutaraldehyde (Electron Microscopy Services, USA) for 10 minutes. Subsequently, negative staining was performed by incubating with 2.5% gadolinium triacetate (Electron Microscopy Services, USA) for 2 minutes. The samples were then viewed using the FEI Tecnai 12 TEM, operating at 100 kV (FEI, USA)^9^.

#### 2.1.8 Evaluation of Dynamin-1 protein concentration in plasma EVs

The protein concentration of DNM1 and Flotillin-1 were quantified using commercially available enzyme-linked immunosorbent assay (ELISA) kits (ABclonal RK01275 and FineTest EH1351 respectively, USA)^16^. The DNM1 protein concentration in the plasma EVs were evaluated by normalizing DNM1 against Flotillin-1 levels. DNM1 levels are presented as pg/ng, which reports the amount of DNM1 in picograms per nanogram of Flotillin-1.

#### 2.1.9 Sample selection

For this analysis, we focused on participants who completed all three questionnaires and blood draws in at least two assessment time points.

#### 2.1.10 Statistical analysis

This analysis was designed to replicate our previous study^9^. First, characteristics between cancer and non-cancer groups were compared using Chi-square test or Fisher’s exact test for categorical variables, and t-test or Mann-Whitney U test for continuous variables. For univariate analysis, Pearson’s correlation test was conducted to evaluate the relationship between the changes in DNM1 expression and FACT-Cog score from baseline. Mann-Whitney U test was also performed to evaluate for differential DNM1 expression (calculated as change in DNM1 levels from baseline) by PCD statuses within each cancer or non-cancer groups. For multivariate analysis, change in DNM1 expression were estimated for each of the four subgroups (Cancer ± PCD, Non-cancer ± PCD) with generalizing estimating equation models (GEE) with a sandwich variance estimator, gaussian family, identity link function and an exchangeable correlation matrix (multivariate and longitudinal analysis). Confounders include baseline DNM1 levels, cancer status, age, gender, marital status, education years, and changes in psychological distress and fatigue. PCD status was designed to be a time-varying variable (e.g. participants with PCD at T2 may not have PCD in T3) due to known heterogeneous trajectories of cognitive function among cancer survivors^17^. All observations were included to facilitate the precise estimation of coefficients. Key coefficients (change/day in DNM1 levels for each subgroup) were calculated based on linear combinations:

- Non-cancer without PCD: time (in days from baseline).
- Non-cancer with PCD: time, PCD status × time.
- Cancer without PCD: time, cancer × time.
- Cancer with PCD: time, cancer × time, PCD status × time, cancer × PCD status × time.

Analyses were tested at a 5% significance level and completed on Stata version 16.1 (College Station, Texas, USA).

### 2.2 Animal Study

#### 2.2.1 Mouse model

All experimental procedures followed NIH vertebrate animal use guidelines and were approved by the UCI Institutional Animal Care and Use Committee (AUP-22-144). 4 months old C57BL/6 female mice (Jackson Laboratory) were housed and maintained on a 12:12 hour light:dark cycle with free access to food and water ad libitum.

#### 2.2.2 Breast cancer induction

Py230-luc (luciferase expressing) murine breast cancer cells were a kind gift from Dr. Ainhoa Mielgo, University of Liverpool, UK. Cells were incubated as a monolayer at 37°C with 5% CO2 in an incubator (Thermo Fisher Scientific) in the F-12K medium (Thermo Fisher Scientific, USA) supplemented with 10% fetal bovine serum (FBS, Gibco), and 0.1% MITO+ serum extender (Corning) and 15% Penn-Strep (Gibco). On the day of tumor implantation, Py230-luc cells were harvested as recommended (ATCC) and prepared as 5×10^5^ cells in a 50 µl hibernation buffer (Hibernate-A medium, Gibco) and 20% polyethylene glycol (PEG, Thermo Fisher Scientific). Tumor cells were implanted in the third mammary gland of each mouse. Hairs were removed using a body cream hair remover (Nair). Py230 cells or buffer were injected directly into the mammary fat pad through a tweezer-uplifted nipple using a 26-gauge needle. The day following injections the mice received 100% aloe vera (Fruit of the Earth) on the injection site to soothe the skin. Animals were monitored closely for one-week post-injection. We did not observe any adverse reaction, and the injection site was completely healed within 2-3 days. Control mice received sham injection treatment using hibernation buffer as above. We used two methods to measure the tumor growth: digital caliper and bioluminescence imaging (BLI). First, the tumor growth was measured using a digital caliper, and tumor volumes were calculated as ellipsoid volume (1/6 π × L × W × (L + W)/2) weekly over eight weeks as described^18^. Concurrently, *in vivo* tumor detection and volume measurements were facilitated by BLI following luciferin injection (GoldBio). Briefly, animals were injected with 0.2 ml (IP) D-luciferin sodium salt (15 mg/ml in PBS). Mice were anesthetized using 2.5% v/v inhalant isoflurane gas anesthesia (VetEquip RC2). Mice resting in a prone position inside the chamber of the imager (IVIS Lumina III, Perkin Elmer) were imaged at a peak emittance time of 12 minutes post-injection. Tumor volumes were reported as relative bioluminescence (photons/s). 1-2 days after the confirmation of tumor growth a cohort of mice was administered a chemotherapy schedule. Mice were treated with an adjuvant chemotherapy regimen using Adriamycin (ADR, doxorubicin hydrochloride, Sigma) and Cyclophosphamide (CYP, Cytoxan, Selleckchem). Mice received ADR made in deionized water at a dose of 2 mg/kg (once weekly, IP) over the course of four weeks. One hour after ADR injections, mice received an injection of CYP made in saline at a dose of 50 mg/kg (once weekly, IP) for four weeks. ADR and CYP were made as injectable solutions at the beginning of the experiment, they were then aliquoted and immediately frozen at −20°C until use. The therapeutic response of tumors to chemotherapy was monitored weekly using the digital caliper and BLI as above. In parallel, noncancer WT female mice received ADR-CYP for comparison of DNM-1 expression with the cancer-bearing mice. At 7-8 weeks post-tumor induction, animals were euthanized by intracardiac perfusion using saline with heparin (10U/mL, Sigma) and 4% paraformaldehyde (Sigma). Fixed brains were immersed in a gradient of sucrose solutions (10 to 30%, Sigma) and sectioned coronally (30 μm thick) using a cryostat (Microm, Epredia). Sections were collected and stored in phosphate-buffered saline (PBS, Gibco) with sodium azide (0.02%) at 4°C.

#### 2.2.3 Dynamin-1 immunofluorescence staining and volumetric analysis

Brain sections were washed with PBS (Gibco) containing 0.3% Tween-20 (Sigma) and then incubated in the 3% hydrogen peroxide (Sigma) solution in PBS and washed with PBS. Sections were then blocked with 10% normal donkey serum (NDS, Jackson Research) in PBS and 0.01% Triton (Sigma) for 45 minutes and then with a primary antibody solution in PBS (rabbit anti-DNM1, 1:250; Synaptic Systems) overnight at 4°C on a see-saw shaker (Enviro-Genie Scientific). Subsequently, sections were washed with PBS and incubated with donkey anti-rabbit Alexa Fluor 568 secondary antibody (Abcam, 1:200) made in PBS with 3% NDS for 1 hour. Sections were washed with PBS, nuclear counterstained using DAPI (15 μM in PBS, Invitrogen), washed again in PBS and mounted on super frost slides using the Slow Fade Antifade Gold mounting medium (Invitrogen).

The immunofluorescent sections were imaged using a laser-scanning confocal microscope (Nikon AX, Japan) equipped with a 40× oil-immersion objective lens (Nikon Plan Apochromat Lambda D, Japan). High-resolution z stacks were acquired using a Nikon Elements AR module at 0.5 μm intervals through the brain section. Unbiased deconvolution for the fluorescent z stacks and in silico volumetric analyses was carried out using our analytical methods as described^19^. The fluorescence signal was deconvoluted to resolve the optimal fluorescence signal at 568 nm wavelength using an adaptive, blinded 3D deconvolution method (ClearView, Imaris v10, Andor Technologies). A 3D algorithm-based surface module (Imaris v10, Andor Technologies) was utilized to evaluate the deconvoluted IMS format images for the volumetric quantification of DNM1 positive immunofluorescent puncta in the hippocampal CA1 and CA3 pyramidal (pyr) and stratum radiatum (sr) subregions. All the Imaris-based data were analyzed using automated batch-processing modules. DNM1 data was expressed as the percentage volume of immunoreactivity to controls.

## 3. RESULTS

### 3.1 Human Study

#### 3.1.1 Descriptive statistics

Samples from 30 cancer and 56 non-cancer participants were eligible. Compared to non-cancer, cancer participants received less education and reported a greater degree of psychological distress (p < 0.05). Most were diagnosed with breast (33%) and head/neck cancers (20%), and underwent systemic treatment with platinum agents (63%) and taxanes (37%). The median DNM1 levels at baseline (pg/ng of Flotillin-1) was significantly lower among cancer participants (29.7 vs 46.1, p = 0.008). This difference remained statistically significant after linear regression adjusted for age, gender, marital status, and education years (β = −15.4, p = 0.023). At T2, there were 7 (23%) cancer who were with PCD. At T3, there were 8 (27%) cancer and 10 (18%) non-cancer participants reporting PCD. Four (13%) cancer participants experienced PCD at both T2 and T3 (**Table 1**).

**Table 1:**
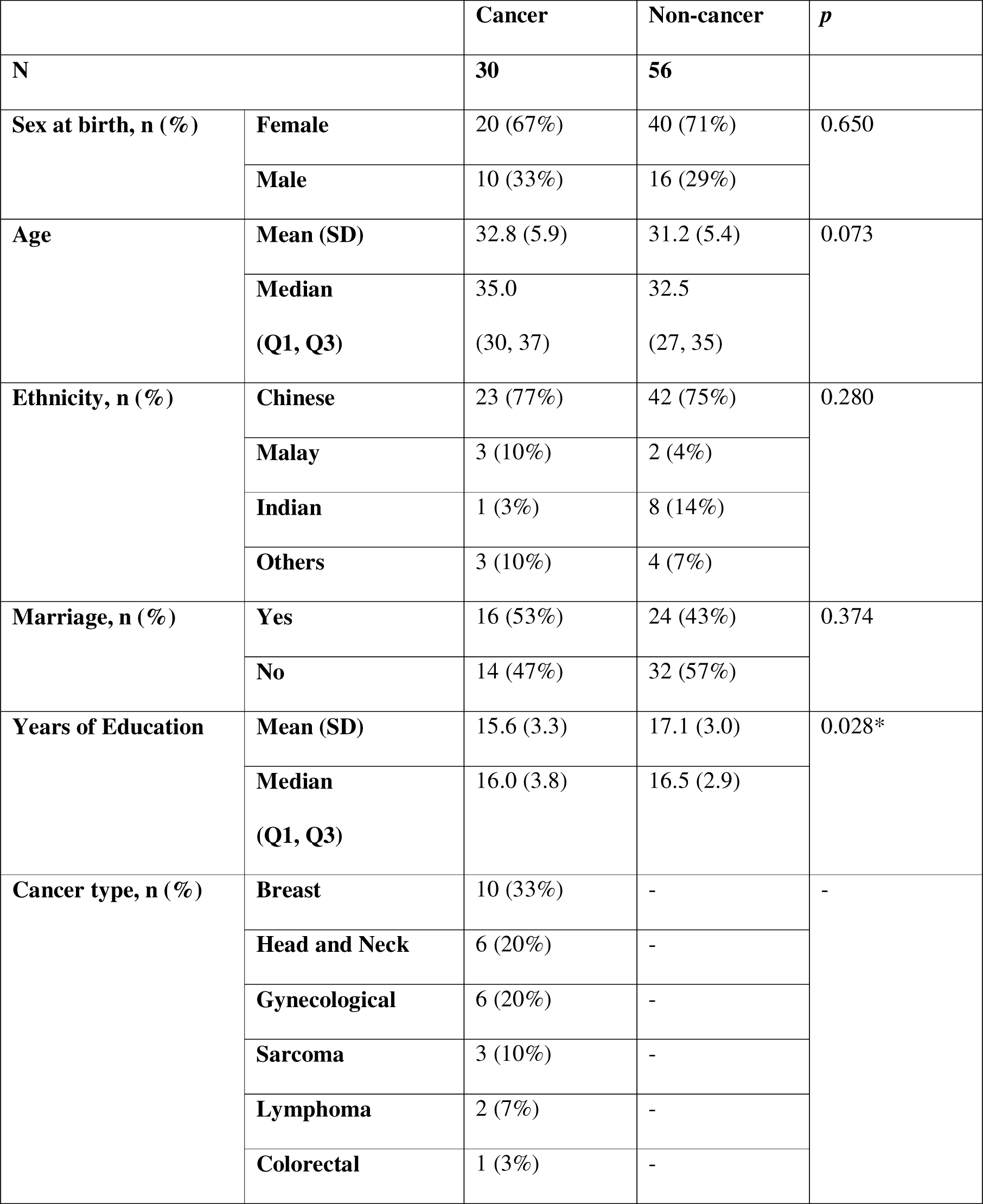
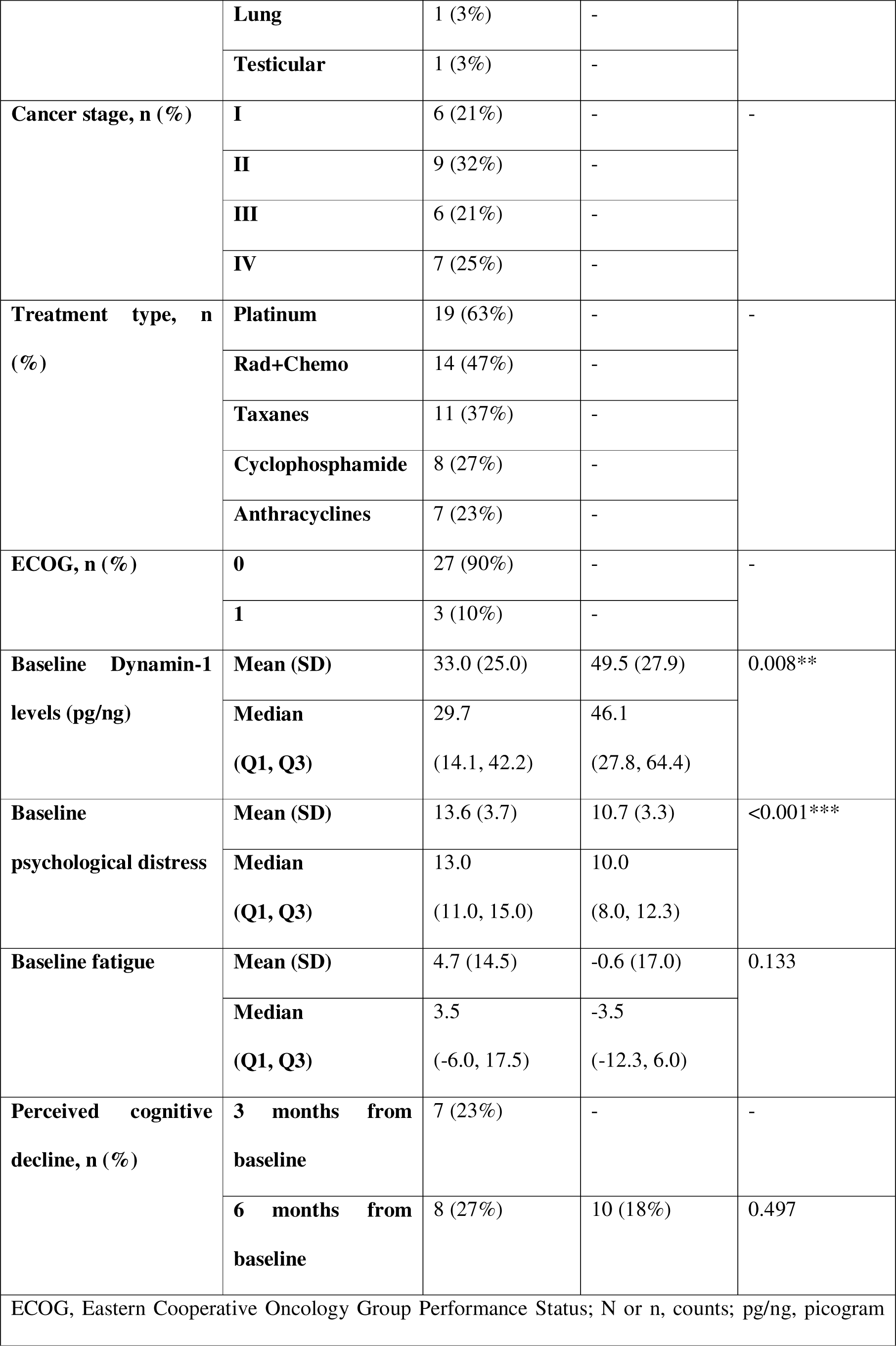
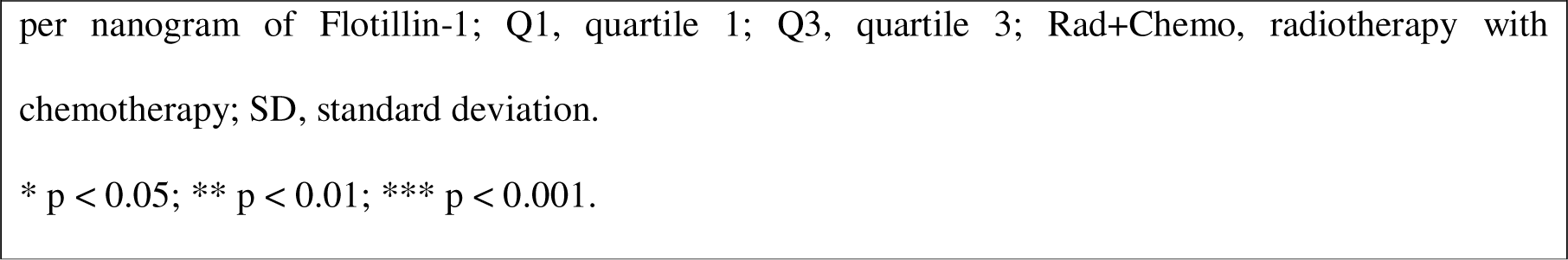
Descriptive statistics comparing cancer with non-cancer participants.

#### 3.1.2 Extracellular vesicle characterization

Plasma EVs displayed spherical morphology (**Fig. 1A-B**) and the presence of the EV marker Flotillin-1. These findings collectively validate the characterization of these plasma EVs.

**Figure 1.**
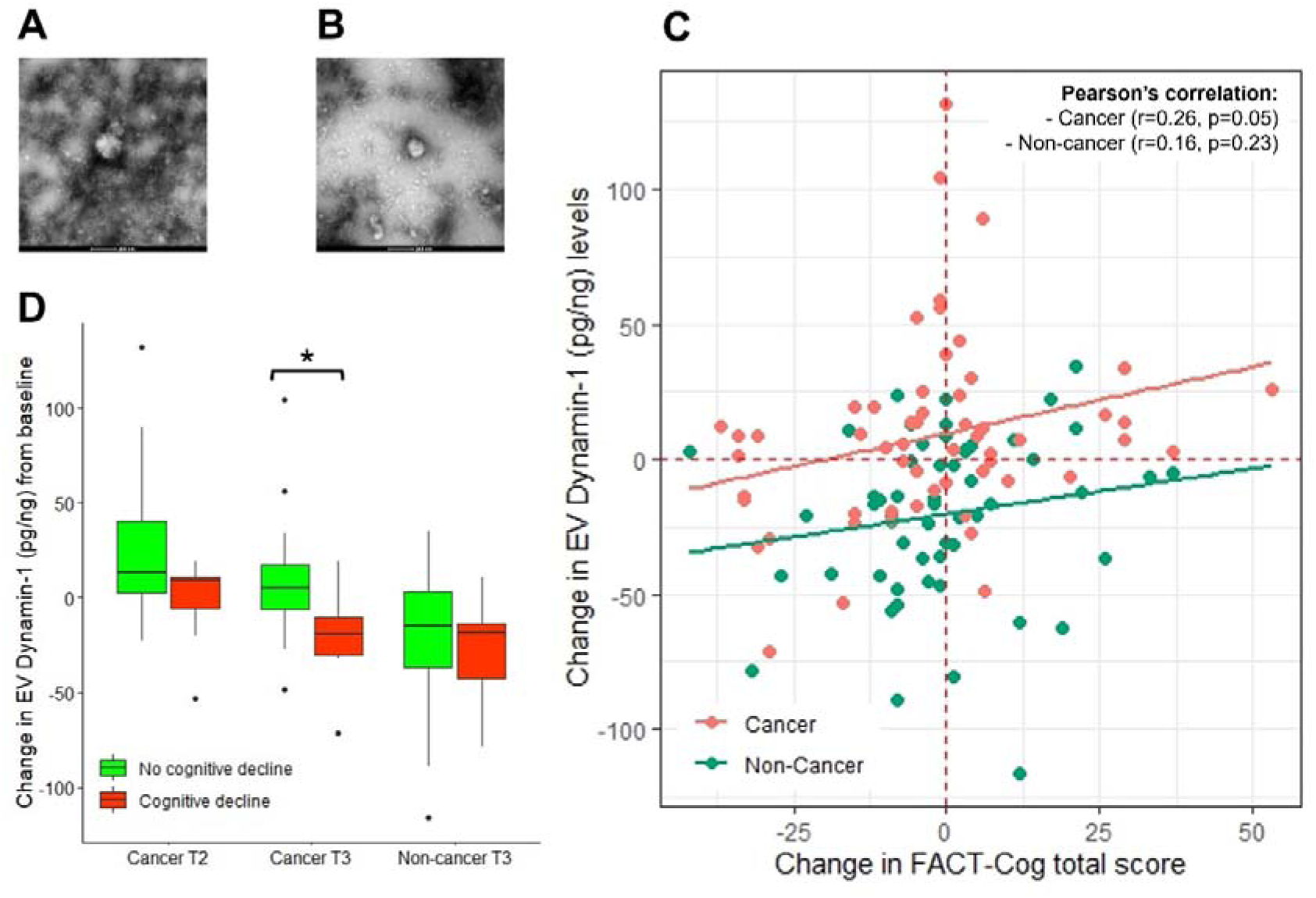
Extracellular vesicle characterization and differential Dynamin-1 expression from human plasma samples stratified by cancer and perceived cognitive decline statuses. A representative electron micrograph of isolated human plasma vesicles of **(A)** cancer and **(B)** non-cancer participants showing the spherical morphology of EVs. Scale bars, 200□nm **(A-B)**. **(C)** Scatter plot correlation between the changes in Dynamin-1 levels (picogram per nanogram of EV marker Flotillin-1) and FACT-Cog total score from baseline. **(D)** Changes in Dynamin-1 levels from baseline were stratified by cancer and perceived cognitive decline statuses, and presented as boxplots (median, interquartile ranges, as well as minimum and maximum excluding outliers). T2 represents the timepoint 3 months after baseline (prior to treatment initiation for cancer participants) while T3 is at 6 months after baseline. * p < 0.05 (D).

#### 3.1.3 Univariate analysis

The Pearson’s correlation coefficient, between changes in DNM1 and FACT-Cog total scores, was statistically significant among cancer participants (r=0.26, p=0.05) but not non-cancer controls (r=0.16, p=0.23) (**Fig. 1C**). At both T2 and T3 among cancer participants, we found a trend of DNM1 downregulation among participants with PCD contrasting the upregulation observed among those without PCD. Statistical significance, when testing for differences between PCD and no PCD, was observed at T3 among cancer participants (p = 0.021). On the other hand, differential DNM1 expression by PCD status was not observed among non-cancer participants (**Fig. 1D**).

#### 3.1.4 Multivariate analysis

By adjusting for confounders with GEE, we found a statistically significant reduction in DNM1 concentration within plasma EVs among cancer participants reporting PCD (β = −0.20, 95% CI = −0.35 to −0.05, p = 0.010) but not in other subgroups (**Table 2**). Characteristics significantly associated with lower DNM1 include age (β = 1.10, 95% CI = 0.14 to 2.08, p = 0.25), Malay ethnicity (β = −17.8, 95% CI = −34.0 to −1.6, p = 0.031) and increased fatigue (β = −0.57, 95% CI = −0.94 to −0.20, p = 0.002).

**Table 2:**
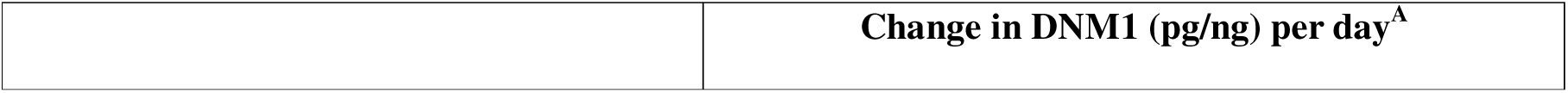
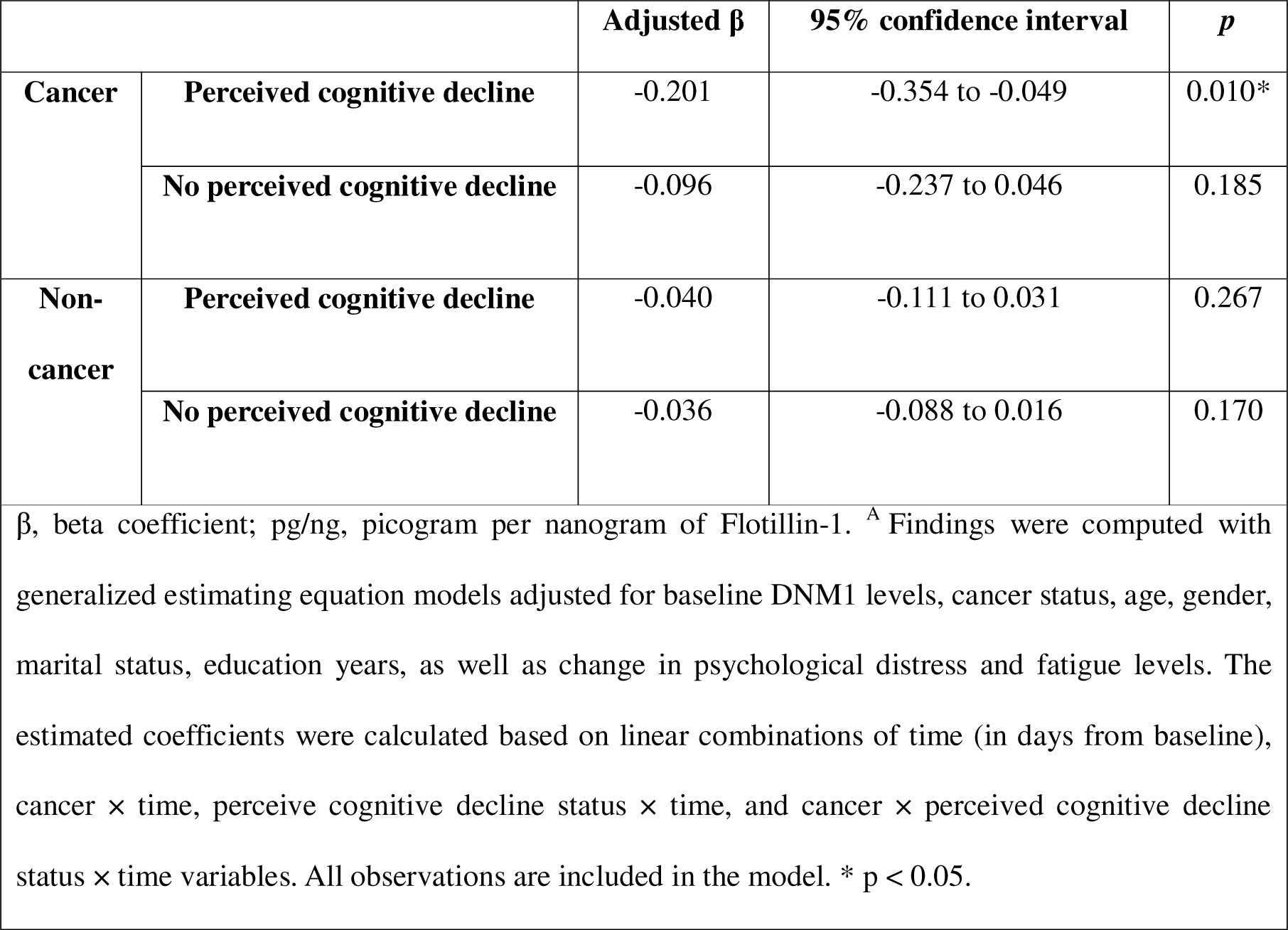
Estimated change in DNM1 levels per day with generalized estimating equation with confounder adjustment.

|  | Change in DNM1 (pg/ng) per day<sup>A</sup> |
| --- | --- |

| | | Adjusted $\beta$ | 95% confidence interval | *p* |
| --- | --- | --- | --- | --- |
| **Cancer** | **Perceived cognitive decline** | -0.201 | -0.354 to -0.049 | 0.010* |
|  | **No perceived cognitive decline** | -0.096 | -0.237 to 0.046 | 0.185 |
| **Non-cancer** | **Perceived cognitive decline** | -0.040 | -0.111 to 0.031 | 0.267 |
|  | **No perceived cognitive decline** | -0.036 | -0.088 to 0.016 | 0.170 |

$\beta$ , beta coefficient; pg/ng, picogram per nanogram of Flotillin-1. <sup>A</sup> Findings were computed with generalized estimating equation models adjusted for baseline DNM1 levels, cancer status, age, gender, marital status, education years, as well as change in psychological distress and fatigue levels. The estimated coefficients were calculated based on linear combinations of time (in days from baseline), cancer × time, perceive cognitive decline status × time, and cancer × perceived cognitive decline status × time variables. All observations are included in the model. \* $p < 0.05$ .

### 3.2 Animal Study

#### 3.2.1 Dynamin-1 expression reduced in the breast cancer-bearing mouse brain treated with adjuvant chemotherapy

Adult WT female mice received an orthotopic injection of murine breast cancer cells (Py230-luc) in the third mammary gland fat pad **(Fig. 2A)**. Mice developed tumors, as indicated by the steady increase in the tumor volume (**Fig. 2B-D**) measured by the BLI and a digital caliper. Both methods of tumor measurements showed comparable tumor growth for the Py230 + Vehicle group **(Fig. 2C-D)**. On the other hand, Py230 breast cancer-bearing mice receiving the adjuvant chemotherapy (ADR-CYP) starting from 10 days post-tumor induction showed significant reductions in tumor volume **(Fig. 2B-D)** indicating a therapeutic response of the tumor to the chemotherapy. Thus, this experimental design provides a testable breast cancer chemobrain model. At 8 weeks post-tumor induction, mice were euthanized, and fixed brains were collected for the immunofluorescence staining and volumetric immunoreactivity quantification of DNM1 (**Fig. 2E-H**). Noncancer WT female mice treated with ADR-CYP were also included for comparison of DNM-1 expression with the cancer-bearing mice. DNM1 expression was observed throughout the hippocampus in the control brains in the hippocampal CA1 and CA3 subregions. Particularly, DNM1 positive puncta were located within the pyramidal layer neuronal soma that emanated through the stratum radiatum (**Fig. 2E** and **2H**). DNM1 positive immunoreactive puncta were quantified using a 3D algorithm-based surface rendering and volumetric quantification. In comparison with the control (no cancer) brains, breast cancer-bearing mice treated with vehicle (Py230-Veh), non-cancer mice treated with chemotherapy (ADR-CYP group), and breast cancer mice treated with ADR-CYP (Py230 + ADR-CYP) had about 13%, 20% and 43% reduction respectively in the DNM1 immunoreactivity in the CA1 region (P<0.04, 0.001, and 0.0001, respectively, **Fig. 2E**). For the CA3 subregion **(Fig. 2G)**, Py230 + ADR-CYP group had a most significant decline in the DNM1 immunoreactivity compared to Controls (P<0.0001, 40% reduction), Py230-Veh (P<0.001, 33% reduction), and noncancer ADR-CYP (P<0.01, 26% reduction) mice brains. In total, breast cancer-bearing mice treated with adjuvant ADR-CYP showed the most drastic reductions in the DNM1 immunoreactivity in the hippocampal CA1 and CA3 subregions important for learning and memory functions.

**Figure 2.**
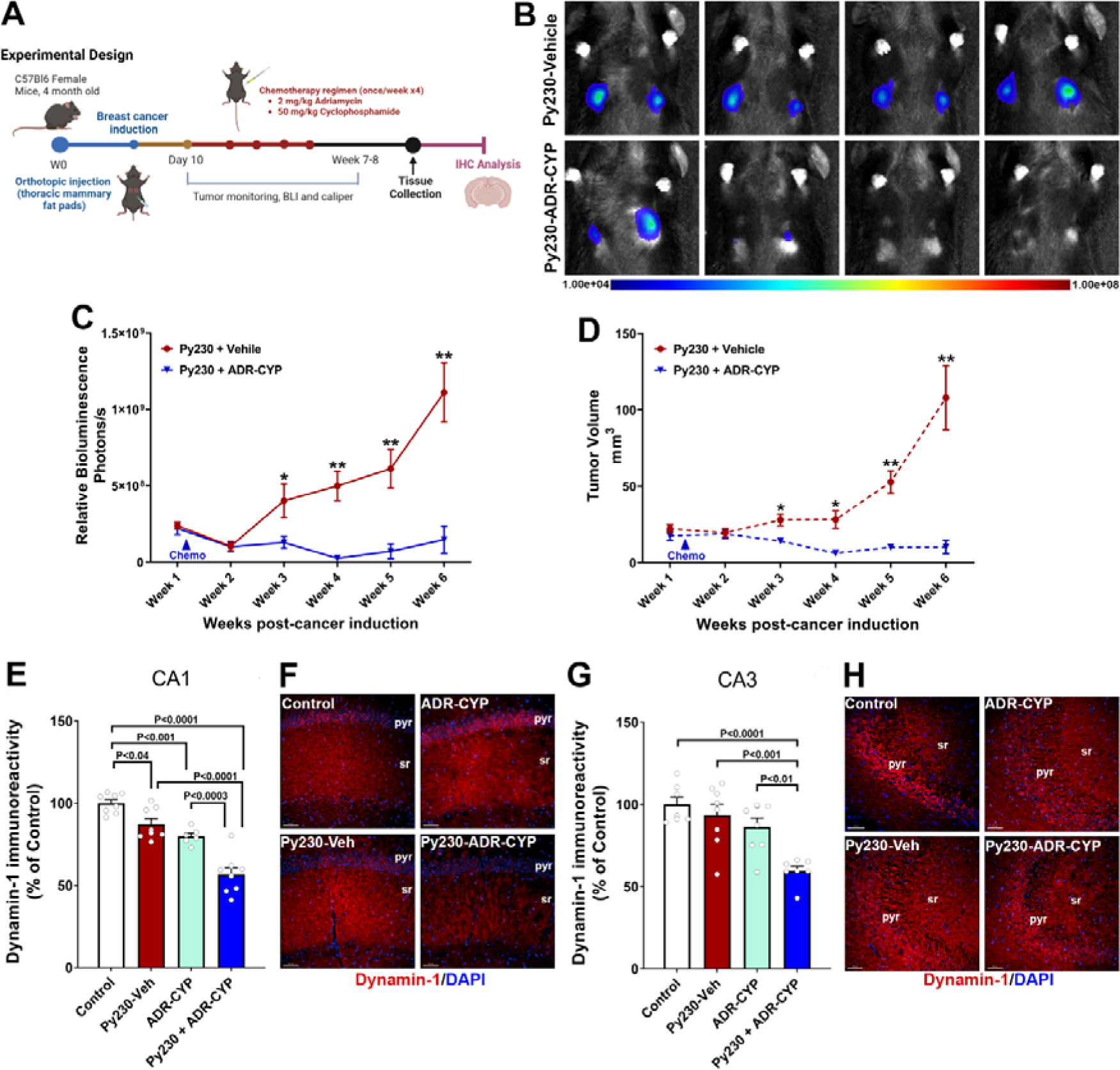
Reduced brain Dynamin-1 in a mouse breast cancer chemobrain model. (A) Animal experimental design: Adult WT female mice received an orthotopic injection of murine Py230-luc breast cancer cells or vehicle in the mammary fat pads, and tumor growth was measured using bioluminescence imaging (BLI) and a digital caliper for 6 weeks post-injection. At 10 days post-tumor induction, a cohort of breast cancer-bearing mice received adjuvant chemotherapy including Adriamycin (ADR, doxorubicin), 2 mg/kg, and cyclophosphamide (CYP), 50 mg/kg, one hour apart, once weekly for 4 weeks. Animals were euthanized and brains were collected at 7-8 weeks post-tumor induction. **(B-D)** Vehicle-treated breast cancer-bearing mice showed a steady increase in the tumor volume as detected by the increased signal *in vivo* **(B)** measured as a relative bioluminescence (Photons/s, **C**). Concurrently, caliper measurements of tumor sites showed an increase in the ellipsoid volume (mm^3^, **D**). Adjuvant chemotherapy (ADR-CYP) significantly reduced tumor volume and growth *in vivo* **(B-D)** thus presenting a breast cancer chemobrain model. **(E-H)** Immunofluorescence staining for Dynamin-1 (red, DAPI nuclear counter stain, blue) showed reduced immunoreactivity in the noncancer (ADR-CYP) and cancer-bearing mice (Py230-Veh, Py230 + ADR-CYP) receiving adjuvant chemotherapy at 8 weeks post-cancer induction compared to controls in the hippocampal CA1 and CA3 pyramidal layer (pyr) and stratum radiatum (sr). Breast cancer-bearing mice treated with ADR-CYP showed most drastic decline in the dynamin-1 immunoreactivity in CA1 and CA3 subregions. Data are presented as mean ± SEM (N = 6-8 mice per group). *P<0.05, and **P<0.01 compared to Py230 + ADR-CYP **(C-D)**. P values were derived from a two-way ANOVA and Tukey’s multiple comparisons test. Scale bars, 50 μm **(F, H)**.

## 4. DISCUSSION

In both humans and animals, we consistently observed that DNM1 downregulation could underlie CRCI pathogenesis. These findings are important and innovative in several ways. We are the first to quantify DNM1 activity within peripheral blood in human samples, and for adding to existing literature regarding the potential of EVs as biomarkers to research on cancer-related complications. Our human study was congruent with our previous study^9^, with reduced DNM1 expressions among cancer patients with perceived cognitive decline following exposure to anticancer treatment. Additionally, we have found a reduction of DNM1 expression in the learning and memory center of the brain, the hippocampal CA1 and CA3 subregions, post-tumor induction and chemotherapy exposure within the animal model, suggesting the potential impact on cognitive function post-cancer. Overall, this study provides important evidence that DNM1 is a biomarker associated with CRCI, and potentially a therapeutic target.

DNM1 plays a crucial role in ensuring effective neurotransmission by facilitating synaptic vesicle recycling via endocytosis, thus DNM1 depletion may reflect reduced neuronal activity dependent on synaptic vesicles^20,21^. DNM1 inhibition reduces hippocampal long-term potentiation and synaptic plasticity, suggesting its importance for the formation of associative memory in the hippocampus^20,22^. DNM1 also plays an important role in neurite development. Previous rodent studies have observed that decreased DNM1 expression cause defects in the biogenesis and endocytic recycling of synaptic vesicles, which impact the neuronal ability to regulate synaptic transmission^23^. Reduction of hippocampal DNM1 in aged mice was related to a decline in hippocampal-dependent memory^24^. There is literature suggesting that expression of DNM1 is associated with genetic polymorphism, with some rare neurological diseases found to be caused by DNM1 mutations^25^. Our previous pre-clinical studies have shown that in addition to reduced neurogenesis^26^, chemotherapy had a detrimental impact on the hippocampal dendritic architecture and reductions in spine density. Particularly, chemotherapy significantly reduced immature spines in the hippocampal dentate gyrus and CA1 regions that plays vital roles in learning and memory function^27^. Taken together, DNM1 could be mediating the structural and functional alterations in the neuronal and synaptic landscape in the brain and contribute to CRCI pathogenesis.

Cancer patients require follow-up care for CRCI up to 10 years post-cancer diagnosis^28^, yet there remains a lack of effective pharmacological management possibly because of the poor understanding of the underlying CRCI mechanism^29^. The etiology of CRCI is likely multifactorial – growing literature suggests that reduction of brain derived neurotrophic factors is associated with CRCI, likely due to the reduction of synaptic plasticity as well as neurogenesis^30^. If cancer-and chemotherapy-induced reduction of DNM1 is truly linked with the long-term neurodegenerative consequences culminating into cognitive impairments, replacement of DNM1 may be a viable therapeutic strategy for ameliorating CRCI. In one study, investigators evaluated the effects of catalpol which was isolated from herbal medicine *Rehmannia glutinosa* and evaluated its impact on synaptic plasticity in aged rat models^31^. The investigators observed that catalpol markedly improved the cognitive function of aged male Sprague-Dawley rats and simultaneously increased the expression of synaptic proteins including DNM1, PSD-95, and synaptophysin in the cerebral cortex and hippocampus, respectively^31^. However, a series of systematic preclinical studies are still required to evaluate the therapeutic role of catalpol in mitigating CRCI. These studies should include breast cancer chemobrain models to establish the underlying mechanistic pathways as well as its safety profile.

There are two key considerations when interpreting our human study findings. First, we have not associated objective cognition with DNM1 levels, which is more widely recognized as a key cognitive endpoint relative to subjective cognition^2^. There is, however, growing literature to demonstrate the usefulness of subjective cognitive measures in epidemiological research and clinical settings^32^. Subjective cognition is arguably a more accurate reflection of how cognitive changes can affect daily functioning compared to objective cognition which is often criticized for the lack of ecological validity^32,33^. While subjective and objective cognitive function may not match for individual patients^34–36^, biomarkers of CRCI, such as pro-inflammatory IL-6 and TNFα, as well as neurogenesis factor BDNF have found consistent directional correlations in studies utilizing both types of cognitive measures^12,30,37^. The biological underpinnings of both dimensions of cognition might be more similar than expected. Second, our current cohort of cancer patients is significantly different in demographic (age and sex) and clinical (cancer diagnoses and anticancer treatment received) characteristics compared to our exploratory cohort of breast cancer patients^9^. Due to limited AYA cancer research and the rarity of this population, the cohort was initially established to assess the prevalence and incidence of CRCI in newly diagnosed AYA cancer patients. Eligibility criteria were strategically designed for age-based recruitment, rather than focusing on cancer or treatment types^11^. With the small sample size in the current analysis, we were unable to incorporate the different cancer-related covariates into the model. Regardless, there are multiple merits with our human study; we have incorporated multiple assessment time points and a non-cancer control following recommendations provided by the International Cognition and Cancer Task Force to facilitate modeling of time-varying cognitive outcomes and isolation of cancer-specific mechanisms^10^. The purposeful recruitment of AYA cancer participants helped with minimizing the confounding effects from age-related neurodegenerative diseases often implicated when studying homogenous cancer populations. While some human data were collected during COVID-19 pandemic wherein social isolation and higher stress levels may have impacted DNM1 levels among non-cancer controls, our multivariable model has accounted for numerous factors including psychological distress, fatigue, and time.

Our breast cancer chemobrain animal model data has added new dimensions of preliminary data that validated the downregulation of hippocampal DNM1 levels exposed to chemotherapy. We found the most drastic reductions in the DNM1 immunoreactivity in the breast cancer-bearing mice (Py230) brains treated with adjuvant chemotherapy (ADR-CYP). Admittedly, without cognitive tests, our animal model did not provide a full picture regarding the cognitive effect of cancer and chemotherapy on DNM1, but it provides significant evidence that DNM1 expression patterns differ between cancer and non-cancer populations. Notably, our previous rodent studies^26,27,38,39^ using ADR and CYP monotherapy in mouse and rat models showed significant neurocognitive decline coincident with the loss of neurogenesis, mature neuron dendritic structure and spine loss in noncancer animals indicating lasting neurodegenerative impact of chemotherapy on brain function. We found significant DNM1 reductions in the hippocampal CA1 and CA3 subregions important for learning and memory, and memory consolidation processes. Such reductions in a functional unit of synaptic vesicle physiology could significantly impact neuronal function, and eventually, cognitive function.

Current literature is lacking with regards to the relationship between brain/hippocampal, plasma and plasma EV DNM1 levels. In comparison to analyzing DNM1 directly in plasma, measuring it within EVs provides insights into the role of cell-to-cell communication, cellular processes and signaling pathways in the pathogenesis of CRCI^40^. In addition, as proteins encapsulated in EVs are stable and have a long half-life, EVs are a more preferred medium for quantifying proteomes in long-term stored biosamples^41,42^. Nevertheless, these elements will be incorporated in our future human and animal studies to further validate and understand the role of DNM1 in CRCI.

In conclusion, we found that downregulation of DNM1 is linked with the onset of CRCI and is consistent in both cancer patients and tumor-bearing mice receiving chemotherapy, strengthening our speculation on DNM1 role as a mediator for CRCI. Upcoming research work should include additional transdisciplinary studies to investigate key mechanisms and generate a clear pathway DNM1 modulation of cognitive function in CRCI trajectories. Further investigation on how this relationship may differ between subjective and objective cognitive impairment, and across different cancer and treatment types will also help to better contextualize the role of DNM1 in CRCI pathogenesis. DNM1 might be a biomarker and therapeutic target for CRCI.

## Declaration of competing interests

The authors have declared that no competing interest exists.

## Author contributions

Conceptualized and designed study: AC. Acquired and analyzed data: DQN, CH, TN, SKG, and YQK. Interpreted data, drafted, revised, and finalized manuscript: DQN, CH, TN, SKG, YQK, MMA, and AC. All authors have read and approved the manuscript for publication.

## Acknowledgements

This work was supported by the National Medical Research Council Singapore (Grant number NMRC/CIRG/1471/2017), US National Institutes of Health (NIH) awards R01CA262213 (MMA), and R01CA276212 (AC and MMA). We would also like to thank Ivy Cheng and Claire Wang for their contribution in participant recruitment and data collection for the human study.

## Data availability

The datasets generated during and/or analyzed during the current study are available from the corresponding author on reasonable request.

## Notes

### Competing Interest Statement

The authors have declared no competing interest.

